# The effect of butyrate-supplemented parenteral nutrition on intestinal defence mechanisms and the parenteral nutrition-induced shift in the gut microbiota

**DOI:** 10.1101/443978

**Authors:** Z Jirsova, M Heczkova, H Dankova, H Malinska, P Videnska, H Vespalcova, L Micenkova, L Bartonova, E Sticova, A Lodererova, L. Prefertusová, A Sekerkova, M Cahova

## Abstract

Butyrate produced by the intestinal microbiota is essential for proper functioning of the intestinal immune system. Total dependence on parenteral nutrition (PN) is associated with numerous adverse effects, including severe microbial dysbiosis and loss of important butyrate producers. We hypothesised that a lack of butyrate produced by the gut microbiota may be compensated by its supplementation in PN mixtures. We tested whether *i.v.* butyrate administration would (a) positively modulate intestinal defence mechanisms and (b) counteract PN-induced dysbiosis. Male Wistar rats were randomised to chow, PN, and PN supplemented with 9 mM butyrate (PN+But) for 12 days. Antimicrobial peptides, mucins, tight junction proteins and cytokine expression were assessed by RT-qPCR. T-cell subpopulations in mesenteric lymph nodes (MLN) were analysed by flow cytometry. Microbiota composition was assessed in caecum content. Butyrate supplementation resulted in increased expression of tight junction proteins (*ZO-1, claudin-7, E-cadherin*), antimicrobial peptides (*Defa 8, Rd5, RegIIIγ*) and lysozyme in the ileal mucosa. Butyrate partially alleviated PN-induced intestinal barrier impairment and normalised IL-4, IL-10 and IgA mRNA expression. PN administration was associated with an increase in Tregs in MLN, which was normalised by butyrate. Butyrate increased the total number of CD4+ and decreased a relative amount of CD8+ memory T cells in MLN. Lack of enteral nutrition and PN administration led to a shift in caecal microbiota composition. Butyrate did not reverse the altered expression of most taxa but did influence the abundance of some potentially beneficial/ pathogenic genera, which might contribute to its overall beneficial effect.

## Introduction

Parenteral nutrition (PN) represents a life-saving treatment in patients with intestinal failure. However, PN and/or lack of enteral feeding are often associated with serious adverse effects, including impaired mucosal homeostasis, loss of immune reactivity (1), compromised intestinal barrier function and generalised sepsis (2).

Proper gut barrier function depends on the integrity of physical barriers, i.e. tight junction proteins and adequate mucin production, sufficient production of antimicrobial compounds by Paneth cells and maintaining an optimal balance between immune tolerance to commensal microbiota and the defence against invading pathogens (3). Lack of enteral feeding significantly affects all of these factors. Paneth cells, which are a specialised type of epithelial cell, release a spectrum of antimicrobial compounds when exposed to alloantigens (4). The absence of enteral feeding decreases mRNA and protein expression of typical Paneth cell antimicrobials like lysozyme, cryptidin-4 and secretory phospholipase A2, thus compromising their function (5, 6). The data concerning the effect of PN on the function of Paneth cells are inconclusive, as their antimicrobial functions have been shown to both increase (3) and decrease (7).

Goblet cells (GCs) continuously secrete glycoproteins (mucins) in order to repair and replace the intestinal mucus barrier (8). Until recently, GCs were considered relatively passive players in promoting intestinal homeostasis and the host defence. However, recent reports indicate that GCs are able to sense and respond to danger signals (such as bacterial pathogens) as well as modulate the composition of the gut microbiome by modifying mucin secretion (9). In a piglet model of enteral nutrition deprivation, GC expansion was established within a few days after the start of total or partial PN (10), which might reflect a higher degradation rate of the mucus layer, a lower rate of mucus secretion, or an altered rate of mucin turnover (11). These data indicate that starvation alters mucus dynamics in the small intestine, which may in turn affect the intestinal defence capacity (11, 12).

The gut microbiota has an irreplaceable role in the maturation of mucosal and systemic immunity (13-15). Depending on its composition, it may either promote a tolerogenic state in the intestinal mucosa (16-20) and instigate mechanisms preventing bacterial overgrowth or induce pro-inflammatory status associated with impaired gut barrier function (21). PN itself, together with a lack of enteral feeding, generates a significant shift in microbiota composition. In rodent models, PN and starvation are associated with decreased gut microbiota diversity, the enrichment of potentially pathogenic and inflammation-promoting species, and the depletion of beneficial anaerobes (3, 7, 22). Heneghan (7) hypothesises that the PN-associated shift in the gut microbiota may be part of a causal relationship with attenuated antimicrobial compound production.

As well as interacting directly with the host intestinal and immune cells, the gut microbiota may affect host intestinal homeostasis via fermentation products. Short-chain fatty acids (SCFA) have multiple beneficial effects on performance and intestinal health (23). SCFA are produced by the fermentation of soluble fibre. To target intestinal SCFA production, an often-used treatment is to supplement the diet with prebiotics (dietary fibre), probiotics (mostly *Lactobacillaceae* or *Bifidobacteriaceae*) or a combination of both. Unfortunately, this approach is not applicable to all situations. Particularly PN-dependent patients with short bowel syndrome often exhibit an increased abundance of *Lactobacillaceae* as well as a lack of butyrate producers in the gut. Therefore, prebiotic/probiotic supplementation may result in D-lactate acidosis or *Lactobacillus* sepsis. The alternative to prebiotic/probiotic treatment is the direct administration of butyrate either *per os* or intravenously. To our knowledge, no study has been published on the effect of i.v. butyrate on the microbiota in a PN context. The purpose of this study was to determine whether the supplementation of a nutrition mixture with butyrate (9 mM) in the absence of enteral feeding would affect immune function and gut microbiota composition. In order to examine this hypothesis, we used a rat model of total parenteral nutrition and assessed the effect of *i.v.* butyrate on Paneth cell function, mucin production, intestine-associated immune cells and the gut microbiome.

## Materials and Methods

### Animals and experimental design

Male Wistar rats (Charles River, initial weight 300-325 g) were kept in a temperature-controlled environment under a 12h light/dark cycle. For PN administration, the right jugular vein was cannulated with a Dow Corning Silastic drainage catheter (0.037 inch) as previously described (3). Control animals underwent the same operation. The catheter was flushed daily with TauroLock HEP-100 (TauroPharm GmbH, Waldbüttelbrunn, Germany). After the operation, the rats were housed individually and connected to a perfusion apparatus (Instech, PA, USA), which allows free movement. For the next 48 hours, the rats were given free access to a standard chow diet (SD, SEMED) and provided Plasmalyte (BAXTER Czech, Prague, CZ) via the catheter at increasing rates (initial rate: 1 ml/hr; goal rate: 4 ml/hr) in order to adapt to the increasing fluid load. Two days after the operation, the rats were randomly divided into three groups. Rats in the experimental groups (PN; PN+But) were provided PN (205 kcal. kg^-1^. d^-1^; 10 hrs per day; rate 4 ml. hr^-1^; light period), the composition of which is given in Table S1. In the PN+But group, the PN mixture was supplemented with 9 mM butyrate. PN alone, PN+But or Plasmalyte was administered for 12 days. All experiments were performed in accordance with the Animal Protection Law of the Czech Republic 311/1997 in compliance with the Principles of Laboratory Animal Care (NIH Guide for the Care and Use of Laboratory Animals, 8^th^ edition, 2013) and approved by the Ethical Committee of the Ministry of Health, CR (approval no. 53/2014).

### Histological evaluation

Tissue samples (distal ileum, proximal colon) were fixed in 4% paraformaldehyde, embedded in paraffin blocks and routinely processed. Sections cut at 4-6 μm were stained with haematoxylin/eosin and examined with an Olympus BX41 light microscope.

### Immunohistochemistry

Paraffin sections (4 μm) were deparaffinised in xylene and rehydrated in graded ethanol. Endogenous peroxidase was blocked, with proteinase K digestion (Dako, Glostrup, Denmark) used for antigen retrieval. The primary anti-lysozyme antibody (rabbit polyclonal, Dako, Glostrup, Denmark) was detected using Histofine Simple Stain Rat MAX PO (Nichirei, Japan). Lysozyme staining intensity was assessed by two independent blinded observers (scale 0 to 3), with average scores presented for each group.

### Flow cytometry

Single cell suspensions from mesenteric lymph nodes (MLN) were obtained by gently fragmenting and filtering the tissues through 100 μm cell strainers (Sigma Aldrich), with lymphocytes isolated by centrifugation on Ficoll (ρ = 1.077 g/ml, GE Healthcare). Isolated cells were frozen and stored at −80°C until analysis. Prior to staining, the lymphocytes were thawed and incubated for two hours in RPMI 1640 + 10% FCS, 2mM L-glutamine, 1% Pen/Strep. Panels for both effector and regulatory T cells were stained simultaneously. First, cells were surface-stained using the following anti-rat antibodies: anti-CD45-FITC (OX-1, Thermo Fisher Scientific), anti-CD4-BV-786 (OX-35, BD Biosciences), anti-CD8α-PerCP-e710 (OX-8, Thermo Fisher Scientific), anti-CD62L-PE (OX-85, SONY) and anti-CD45RC-Alexa Fluor 647 (OX-22, SONY) for the effector T-cell panel, and anti-CD45-FITC, anti-CD4-BV-786 and anti-CD25-PE (OX-39, Thermo Fisher Scientific) for the regulatory T-cell panel. Second, the cells in both panels were fixed and permeabilised using an intracellular staining kit (Anti-Mouse/Rat Foxp3 Staining Set APC, Thermo Fisher Scientific) either with Foxp3 antibody (FJK-16s, regulatory T cells) or PBS (effector T cells) in conjunction with 15-min blocking using 2% normal rat serum (regulatory T cells only, Thermo Fisher Scientific). Immediately after staining with anti-Foxp3 mAb, the lymphocytes were analysed using the BD LSR II flow cytometer (BD Biosciences).

### RT-qPCR

Pieces of the distal ileum (5-8 cm from the ileocaecal valve) were rapidly dissected, flushed first with cold saline and then with RNA later, opened along the mesenteric border and the mucosa was then scraped using a glass slide and immediately frozen in liquid nitrogen. To determine cytokine expression, Peyer’s patches were dissected from the rest of the ileum. Total RNA was extracted using the RNeasy PowerMicrobiome Kit (Qiagen, Hilden, Germany). A DNAase step was included to avoid possible DNA contamination. A standard amount of total RNA (1600 ng) was used to synthesise first-strand cDNA with the High Capacity RNA-to-cDNA Kit (Applied Biosystems, Foster City, CA, USA). The RT-PCR amplification mixture (25ul) contained 1 ul template cDNA, SYBR Green Master Mix buffer (QuantiTect, Qiagen, Hilden, Germany) and 400nM (10 pmol/reaction) of sense and antisense primers. Primers were designed based on known rat sequences taken from the GeneBank Graphics database: *https://www.ncbi.nlm.nih.gov*. Primer design was performed with Primer3 software: http://www.frodo.wi.mit.edu (Table 1). The reaction was run on the ViiA 7 Real-Time PCR System (Thermo Fisher Scientific, USA). Results were analysed using SDS software, ver. 2.3 (Applied Biosystems, Foster City, CA, USA). The expression of genes of interest was normalised to the housekeeper gene Rplp2 and calculated using the ΔΔCt method.

**Table 1.**
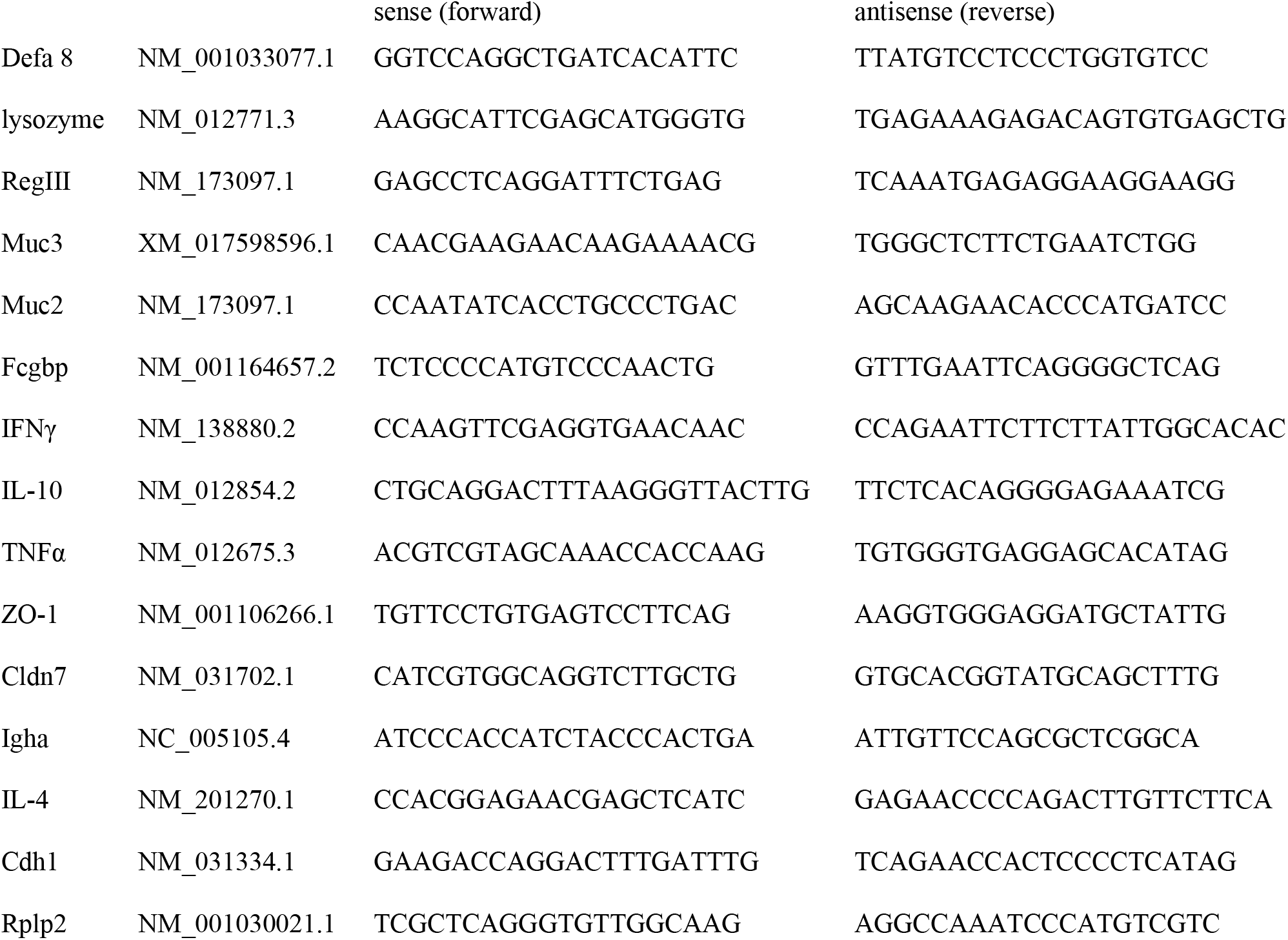
Primer sequences (5’ - 3’).

### Determination of microbiota composition

Microbiota composition was determined in caecum content. All samples were frozen at −20 °C until required. DNA was isolated using the QIAamp PowerFecal DNA Kit (Qiagen). Extracted DNA was used as a template in amplicon PCR to target the hypervariable V4 region of the bacterial 16S rRNA gene. A 16S metagenomics library was prepared according to the Illumina 16S Metagenomic Sequencing Library Preparation protocol, with some modifications described below. Each PCR was performed in triplicate, with the primer pair consisting of Illumina overhang nucleotide sequences, an inner tag and gene-specific sequences (forward: TCGTCGGCAGCGTCAGATGTGTATAAGAGACAG-InnerTag-GTGYCAGCMGCCGCGGTAA; reverse: GTCTCGTGGGCTCGGAGATGTGTATAAGAGACAGC-InnerTag-GGACTACNVGGGTWTCTAAT) (24, 25). The Illumina overhang served to ligate the Illumina index and adapter. Each inner tag – a unique sequence of 7–9 bp – was designed to differentiate samples into groups. After PCR amplification, triplicates were pooled and the amplified PCR products determined by gel electrophoresis. PCR clean-up was performed with Agencourt AMPure XP beads (Beckman Coulter Genomics). Samples with different inner tags were equimolarly pooled based on fluorometrically measured concentrations using the Qubit^®^ dsDNA HS Assay Kit (Invitrogen™, USA) and microplate reader (Synergy Mx, BioTek, USA). Pools were used as a template for the second PCR with Nextera XT indexes (Illumina, USA). Differently indexed samples were quantified using the KAPA Library Quantification Complete Kit (Kapa Biosystems, USA) and equimolarly pooled according to the measured concentration. The prepared library was checked with the 2100 Bioanalyzer Instrument (Agilent Technologies, USA), with concentrations measured by qPCR shortly prior to sequencing. The library was diluted to a final concentration of 8 pM with the addition of 20 % PhiX DNA (Illumina, USA). Sequencing was performed using the Miseq Reagent Kit v2 according to the manufacturer’s instructions (Illumina, USA).

### Data processing and statistical analysis

Sequencing data, i.e. raw sequences, were processed using standard bioinformatic procedures within QIIME 1.9.1 package (26). In short, these include quality filtering, chimera removal, open reference clustering and taxonomic identification based on the SILVA 123 database and UCLUST algorithm (27). Raw sequences were filtered according to default quality requirements in QIIME 1.9.1 (-r: 3; -p: 0.75; -n:0; -q:3). Chimeras were detected and filtered using the UCHIME algorithm with the Gold database. Data were afterwards clustered at the 97% similarity threshold against SILVA database version 123. Representative sequences were aligned, and a phylogenetic tree was constructed and taxonomic identity determined by the USEARCH algorithm. The data were treated as compositional (proportions of total read count in each sample, non-rarefied) and prior to all statistical analyses were transformed using centered log-ratio transformation (28). Sequencing data are available from ENA database under the accession number PRJEB28521. All analyses were performed in R, version 3.4.2. (29).

Gene expression data and flow cytometry data are presented as mean ± SD. Statistical analysis was performed using the Kruskal-Wallis test with multiple comparisons. Differences were considered statistically significant at the level of p<0.05. For testing group pairwise differences in microbial composition, we applied ANOVA test with Tukey’s honest significance. The statistical analyses were performed on each of the six taxonomy levels (Phylum, Class, Order, Family, Genus and OTUs) separately. The resulting p-values were adjusted for multiple hypothesis testing using the Benjamini-Hochberg procedure. Results were considered significant at FDR<=10%. Hierarchical clustering with Euclidean distance and the average-linkage algorithm was used to cluster microbial profiles in the heatmap and the radar chart.

## Results

### Ileal and colonic architecture

Compared with controls, we observed significantly reduced mucosal thickness in the ileum (550±40 vs 746±28 μm, p<0.05) and colon (886±90 vs 2750±110 μm, p<0.01) (Fig 1) in rats totally dependent on PN. Butyrate supplementation had no effect on these parameters (ileum: 535±32; colon: 1020±103 μm).

**Fig 1.**
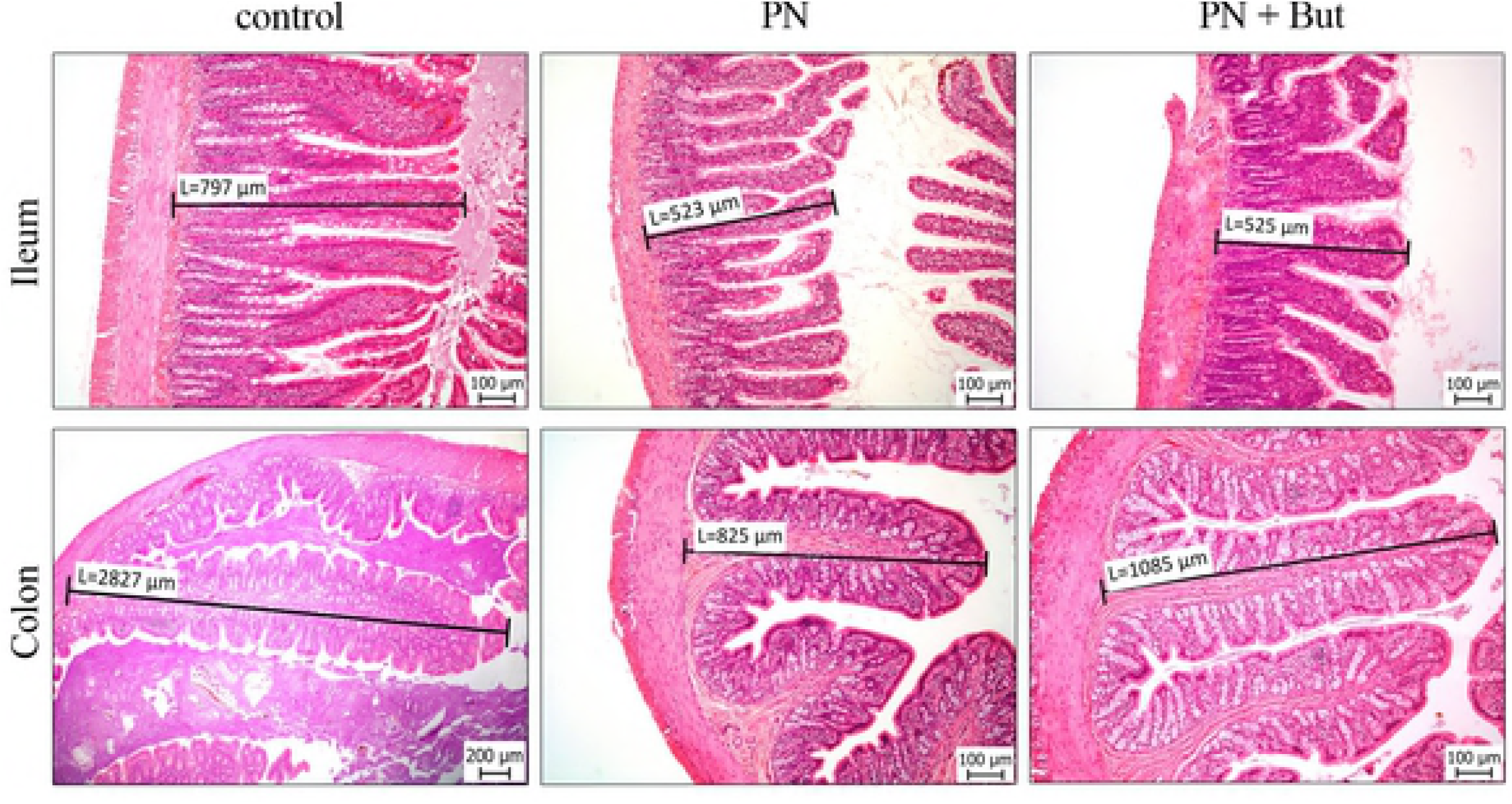
Histology of the intestinal mucosa. Mucosal thickness was assessed in the small intestine (ileum) and the large intestine (colon). Sections of intestinal tissues were stained with H&E (magnification x100).

### Butyrate stimulates Paneth cell function

To examine the potential Paneth cell alterations associated with butyrate administration, we determined the expression of Paneth cell-produced compounds. First, we examined the expression of lysozyme. Immunohistochemical staining confirmed its presence in Paneth cell granules in the ileum in all groups (Fig 2). Based on staining intensity, PN administration substantially increased lysozyme expression compared with controls (Fig 2B). Supplementation of the PN mixture with butyrate resulted in the further elevation of lysozyme-specific staining intensity (Fig 2C). Corresponding results were obtained at the mRNA level (Fig 2E). Next, we determined the expression of other antimicrobial peptides, i.e. α-defensins (Rd5, Defa8) and RegIIIγ (Figs 2F-H). Whereas PN alone had no effect, we found significantly increased expression of all three compounds in the PN+But group. In conclusion, our data show that supplementation of a PN mixture with butyrate is associated with increased Paneth cell function, as measured by the expression of antimicrobial peptides.

**Fig 2.**
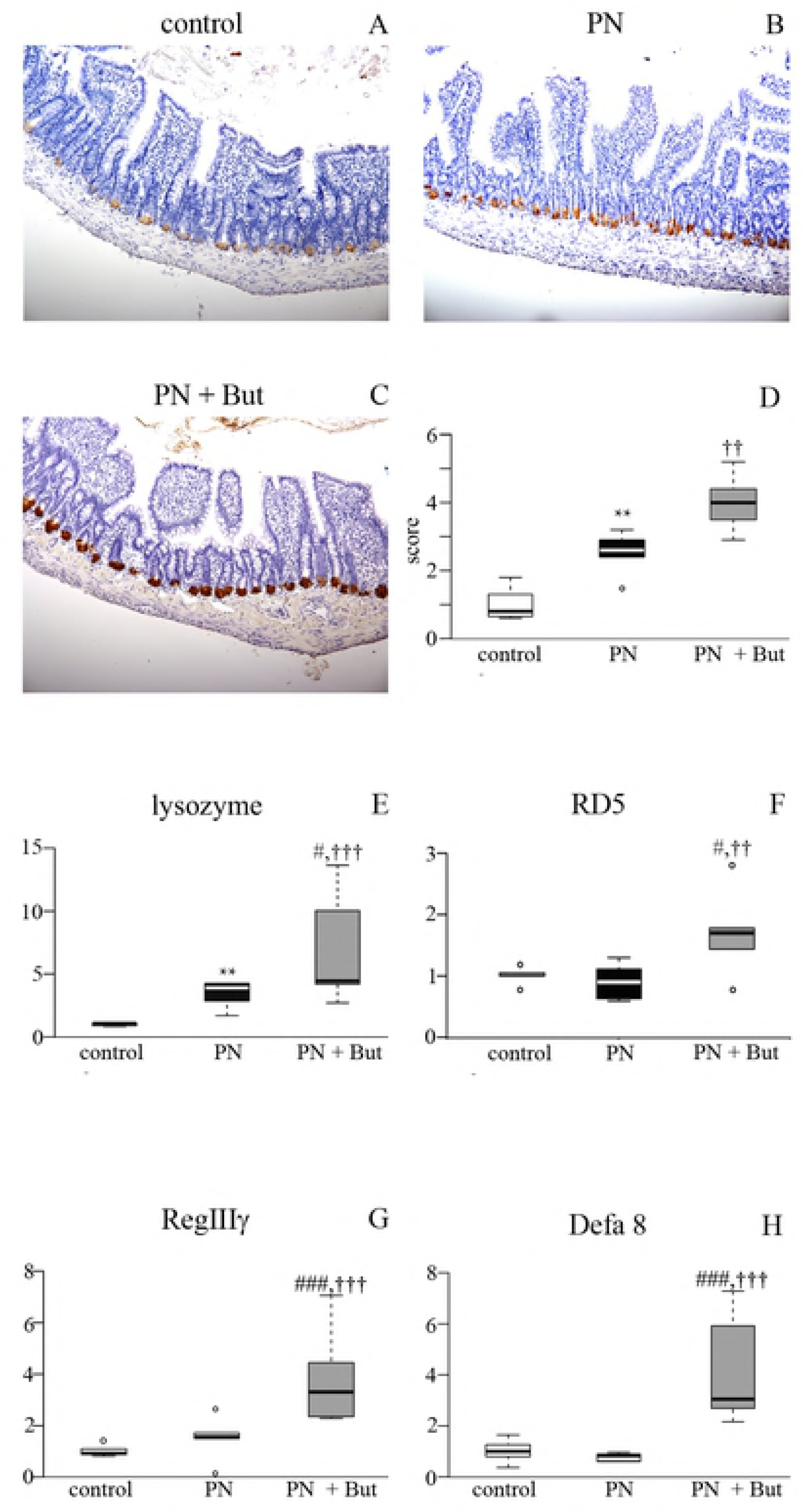
Host defence peptide proteins and mRNA expression in the ileum. A-C: Lysozyme staining, magnification x 200; D: lysozyme staining quantification; E: lysozyme mRNA expression; F: RD5 mRNA expression; G: Defa8 mRNA expression; H: RegIIIγ mRNA expression. mRNA expression is given as a fold change over the control group. Results are presented using Tukey box-and-whisker plots as quartiles (25%, median, and 75%). ^**^p <0.01 PN vs control; ^††^p <0.01; ^†††^p<0.001 PN+But vs control; ^#^p<0.05, ^###^p<0.001 PN+But vs PN.

### Butyrate promotes mucin production

GCs specialise in producing and secreting mucin glycoproteins and other factors to form a protective mucus layer in the intestine. We assessed their function according to the number of GCs (normalised as the GC number per 200 enterocytes) and by mRNA expression of three GC products in the ileum. GC numbers tended to be higher in the PN group compared with controls, but the difference was not statistically significant (53.2±5 vs 44±9.4, p=0.08). The addition of butyrate resulted in a further increase in abundance (63.6±8.5, p<0.01 vs controls, p<0.05 vs PN). Expression of *Muc2*, the main secretory mucin, increased in the PN group compared with controls, and was further potentiated by butyrate. *Muc3*, the dominant transmembrane mucin, was elevated only in the PN+But group. *Fcgbp* expression was not affected in any group (Fig 3). These data indicate that in response to the absence of enteral feeding GCs increase activity, and that butyrate supplementation significantly stimulates this process.

**Fig 3.**
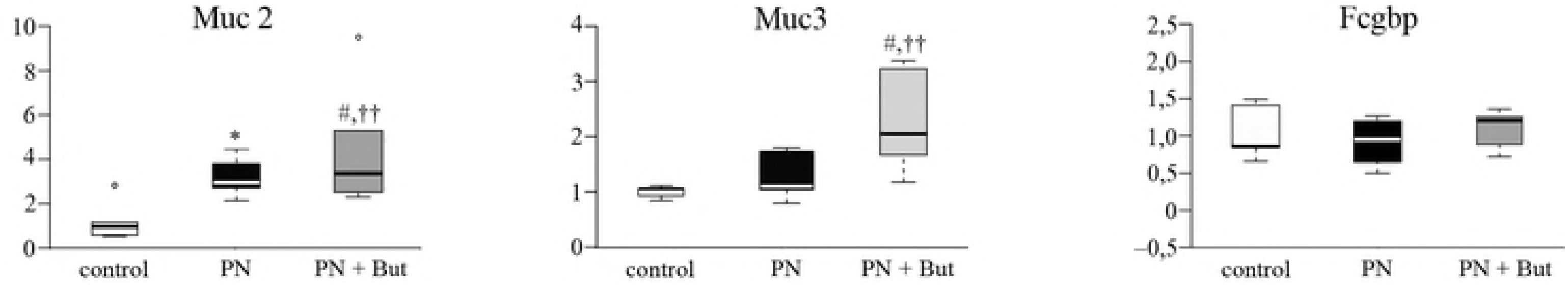
mRNA expression of mucosa-forming genes in the ileum. A: Muc2; B Muc3; C: Fcgbp. mRNA expression is given as a fold change over the control group. Results are presented using Tukey box-and-whisker plots as quartiles (25%, median, and 75%). ^*^p<0.05 PN vs control; ^††^p<0.01 PN+But vs control; ^#^p<0.05; ^##^p<0.01 PN+But vs PN.

### Butyrate alleviates PN-induced small intestinal permeability

The effect of butyrate on small intestinal integrity was assessed by *in vitro* permeability for HRP and by the expression of tight junction proteins. Ileal segments of both the PN and PN+But groups were more permeable for HRP compared with controls (Fig 4A). Butyrate supplementation decreased intestinal permeability compared with the PN group, although it did not match the control level. The expression of tight junction proteins (*ZO-1, claudin-7, E-cadherin*) was similar in the control and PN groups and significantly increased in the PN+But group (Figs 4B-D). In summary, these findings support the hypothesis that butyrate alleviates the detrimental effect of PN on intestinal permeability via the stimulation of tight junction protein expression.

**Fig 4.**
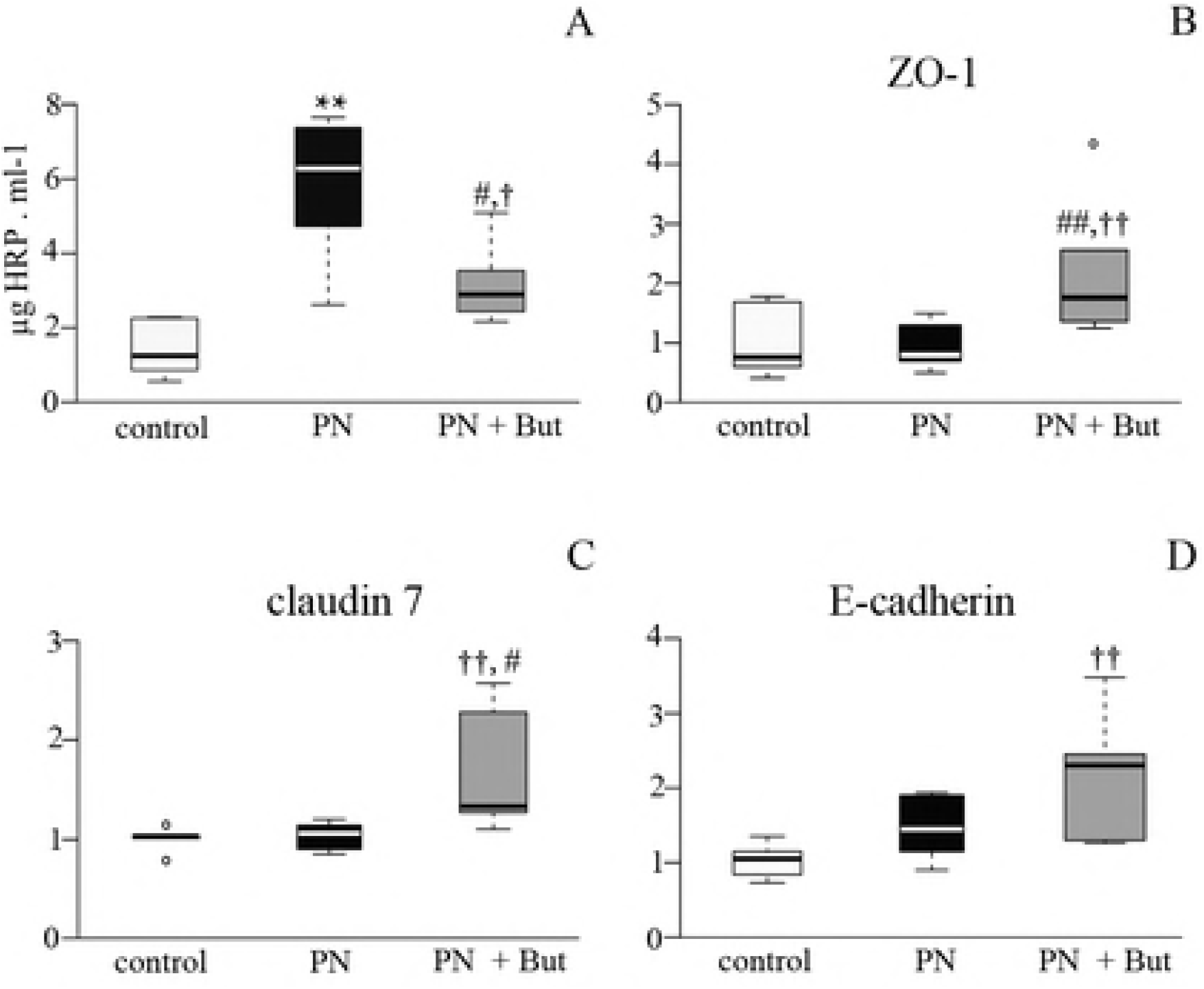
The effect of butyrate on tight junction proteins mRNA expression and intestinal permeability. A: *ZO-1* mRNA; B *E-cadherin* mRNA; C *claudin-7* mRNA; D: HRP leakage *in vitro*. mRNA expression is given as a fold change over the control group. Results are presented using Tukey box-and-whisker plots as quartiles (25%, median, and 75%). ^*^p<0.05 PN vs control; ^††^p<0.01 PN+But vs control; ^#^p<0.05; ^##^p<0.01 PN+But vs PN.

### The effect of butyrate on lymphocyte phenotypes and cytokine expression

In order to determine the effect of butyrate on gut-associated T-cell subpopulations, we isolated lymphocytes from MLN and analysed them by flow cytometry (Fig 5). In MLN, PN alone did not affect the total number of CD4+ or CD8+ lymphocytes, CD4+/CD8+ ratio (2.3±0.5 vs 2.4±0.4) or percentage of different CD8+ subpopulations, but it did increase the percentage of CD4+Foxp3+CD25+ (Treg). Butyrate supplementation led to a significant rise in CD4+ lymphocytes but did not change the total number of CD8+ lymphocytes, resulting in an increased CD4+/CD8+ ratio (3.5±0.2).

**Fig 5.**
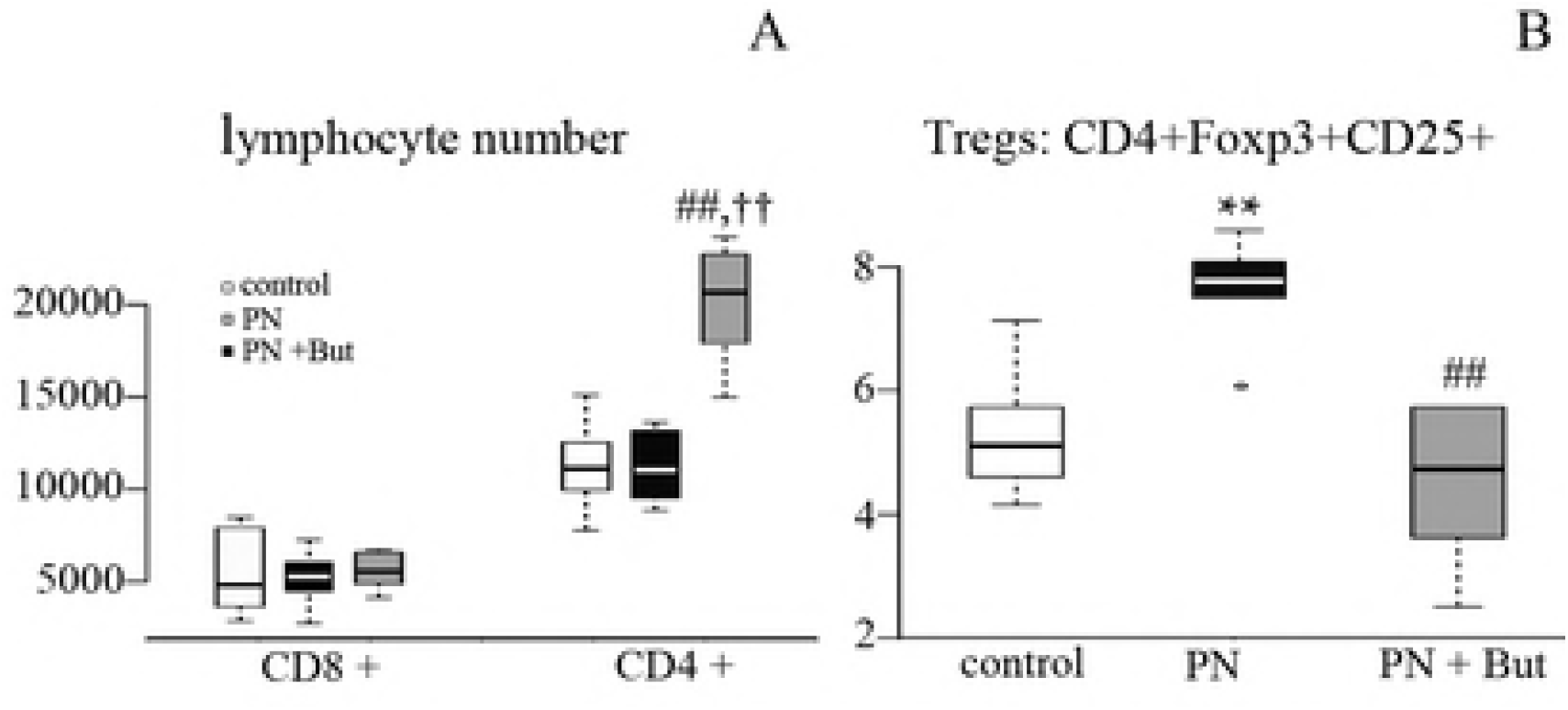
The effect of butyrate on the distribution of T-cell subpopulations in mesenteric lymph nodes. A: Total CD4+ and CD8+ lymphocyte numbers; B: Tregs subpopulation. Results are presented using Tukey box-and-whisker plots as quartiles (25%, median, and 75%). ^**^p<0.01 PN vs control; ^††^p<0.05 PN+But vs control; ^##^p<0.01 PN+But vs PN.

In the PN group, we found significant attenuation of IL-10 (Fig 6A) and IL-4 mRNA (Fig 6B) expression in Peyer’s patches as well as IgA mRNA expression (Fig 6C) in the intestinal mucosa. In the PN+But group, the expression of both cytokines increased to the levels observed in controls and IgA expression was nearly normalised. Taken together, butyrate added to a PN mixture is associated with an increase in the total CD4+ lymphocyte population, normalisation of the Tregs subpopulation in MLN, and an increase in gut mucosal immunity.

**Fig 6.**
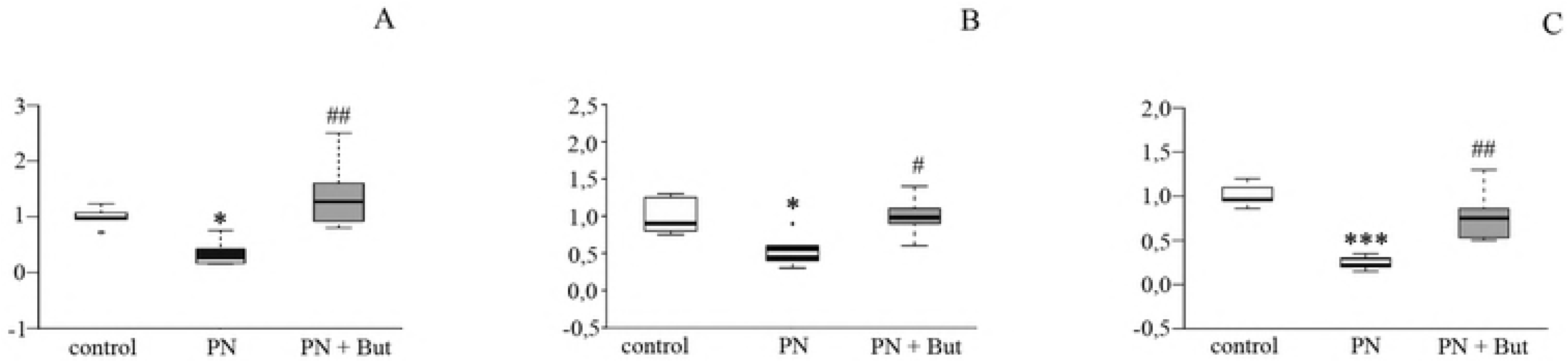
The effect of butyrate on cytokine and IgA mRNA expression. A: IL-10 expression in Peyer’s patches; B: IL-4 expression in Peyer’s patches; C: IgA expression in the intestinal mucosa. mRNA expression is expressed as a fold change over the control group. Results are presented using Tukey box-and-whisker plots as quartiles (25%, median, and 75%). ^*^p<0.05, ^***^p<0.001 PN vs control; ^#^p<0.05; ^##^p<0.01 PN+But vs PN.

### The effect of butyrate supplementation on the microbiota

Microbiota composition was assessed via sequencing of the 16S rRNA gene in caecum content sampled at the time of sacrifice. Alpha diversity was assessed in terms of species richness (OTU numbers, Chao1 index) or evenness (Shannon index, Simpson index) (Table 2). Caecal microbiota in PN+But group tend to be less diverse compared with control or PN groups but this tendency reached the statistical significance only when OTUs number is concerned.

**Table 2.** Alpha diversity.

|  | OTUs | Chao1 | shannon | simpson |
| --- | --- | --- | --- | --- |
| control | 477 (180) | 814 (340) | 6.23 (1.14) | 0.96 (0.03) |
| PN | 429 (109) | 726 (289) | 6.04 (1.43) | 0.96 (0.1) |
| PN+But | 328 (157)<sup>†</sup> | 583 (288) | 4.53 (3.44) | 0.85 (0.34) |

Data are given as median and IQR. <sup>†</sup> $p < 0.05$ PN+But vs control

The absence of enteral feeding in combination with PN administration had a significant effect on gut microbiota composition. At the phylum level, *Proteobacteria* significantly increased in both PN-dependent groups. Butyrate administration was associated with a decrease in *Proteobacteria* abundance but this trend did not reached statistical significance. Butyrate supplementation counteracted the deregulation of *Cyanobacteria* observed in the PN group (Fig 7).

**Fig 7.**
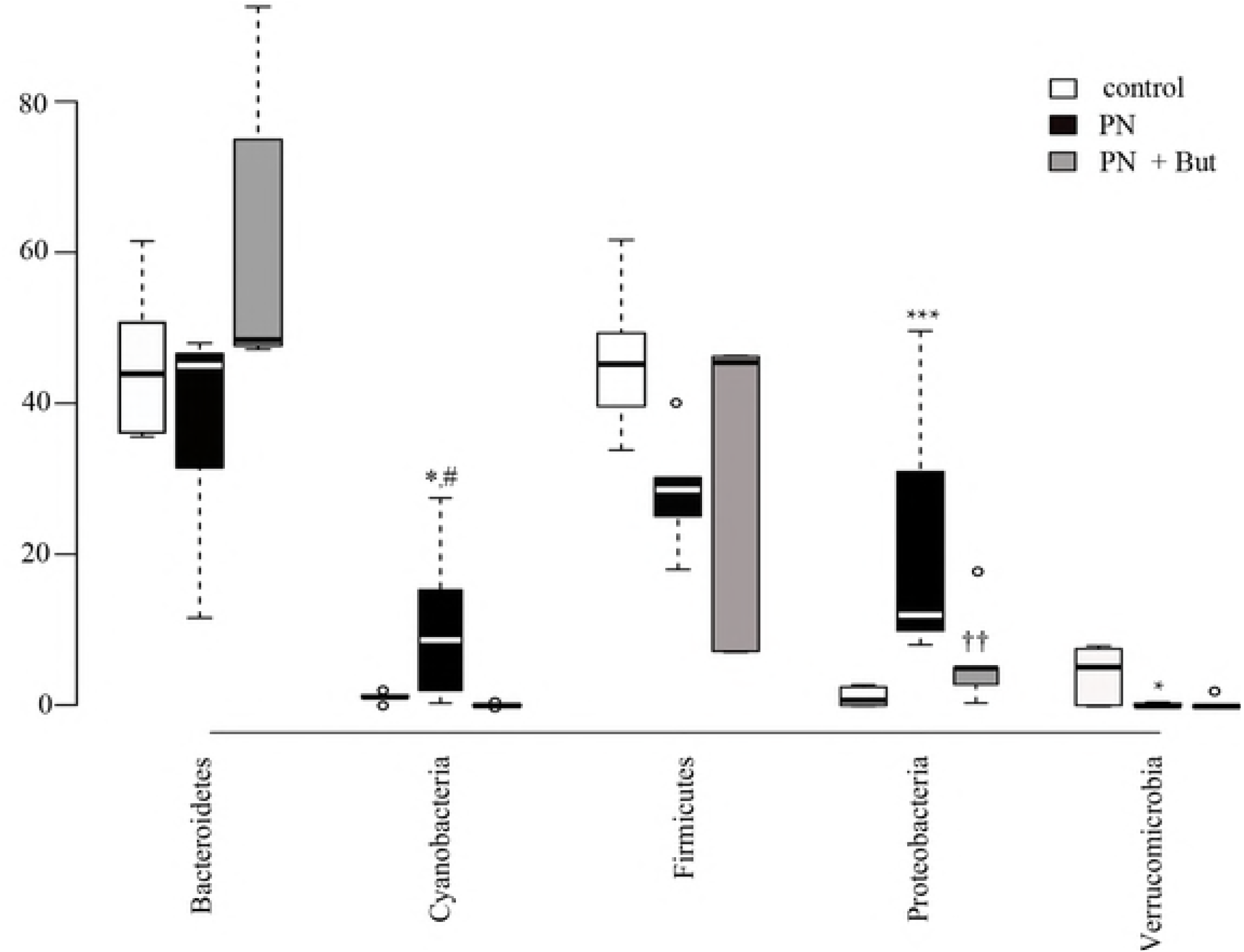
Microbiota composition in the caecum: phylum level. Results are presented using Tukey box-and-whisker plots as quartiles (25%, median, and 75%) and outliers (open circles). ^*^p<0.05, ^***^p<0.001 PN vs control; ^††^p<0.05 PN+But vs control; ^#^p<0.05 PN+But vs PN.

The distribution pattern of abundant (<1%) bacterial families is shown in Fig 8. *Porphyromonadaceae* and *Alcaligenaceae* were significantly elevated while the *Clostridiales vadinBB60* was reduced in both PN-dependent groups compared with controls. The abundance of *Bacteroidaceae, Enterobacteriaceae, Lachnospiraceae,* and *Lactobacillaceae* was significantly altered only in one of the PN-dependent groups compared with controls but the trend was similar, i.e. of the same orientation, in both of them. Butyrate supplementation had significant effect on the abundance of *Peptococcaceae* and one unidentified taxon belonging to *Gastranaerophilales*.

**Fig 8.**
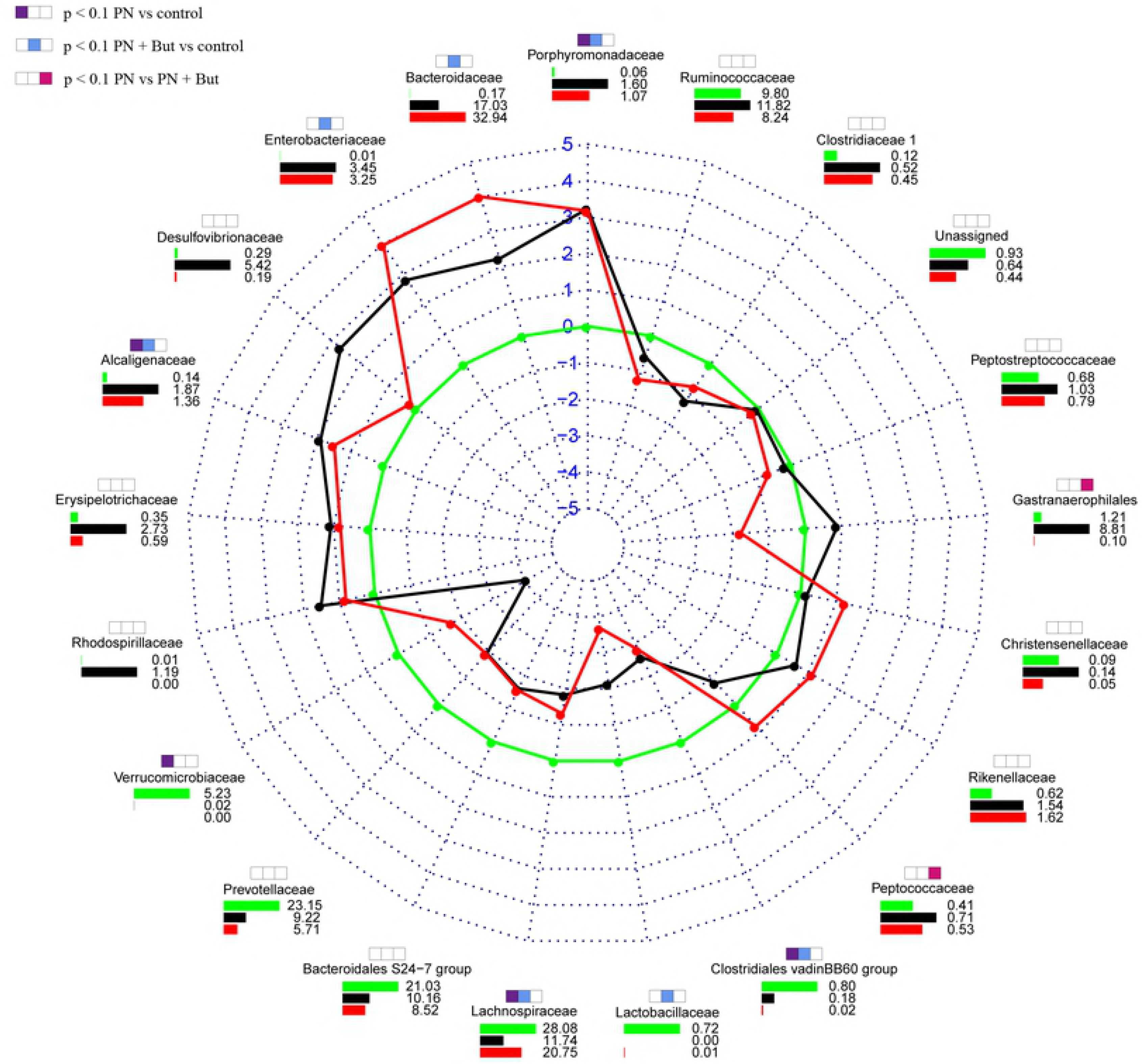
Distribution pattern of all abundant (<1%) bacterial families in the caecum. Lines show the fold change vs controls, green line represents the null change (controls vs controls). Bar charts demonstrate the relative abundance (%) of each family. Colour key: green = controls; black = PN; red = PN+But. The significance (p<0.05) is shown using coloured boxes above the family names.

We identified 20 genera that were significantly differently (p < 0.05) represented in at least on of the PN-dependent group compared with controls (Fig 9). Five genera were deregulated in both the PN and PN+But groups, i.e. *Bacteroides, Parabacteroides, Alistipes, Parasutterella* (increased) and *Prevotellaceae NK4A214 group* (decreased). Compared with the PN group, butyrate supplementation resulted in the increased abundance of *Anaerostipes, Lachnospiraceae AC2044 group* and *Roseburia*, but decreased the representation of the *Prevotellaceae Ga6A1 group* and unidentified bacteria from the *Gastranaerophilales* order. Similar trend (PN+But < PN) was observed in case of *Desulfovibrio sp.* (p=0.055). All relevant statistical data are shown in Table S2. Our data confirm the profound effect of the lack of enteral feeding on microbiota composition. Butyrate supplementation counteracted only some of the alterations.

**Fig 9.**
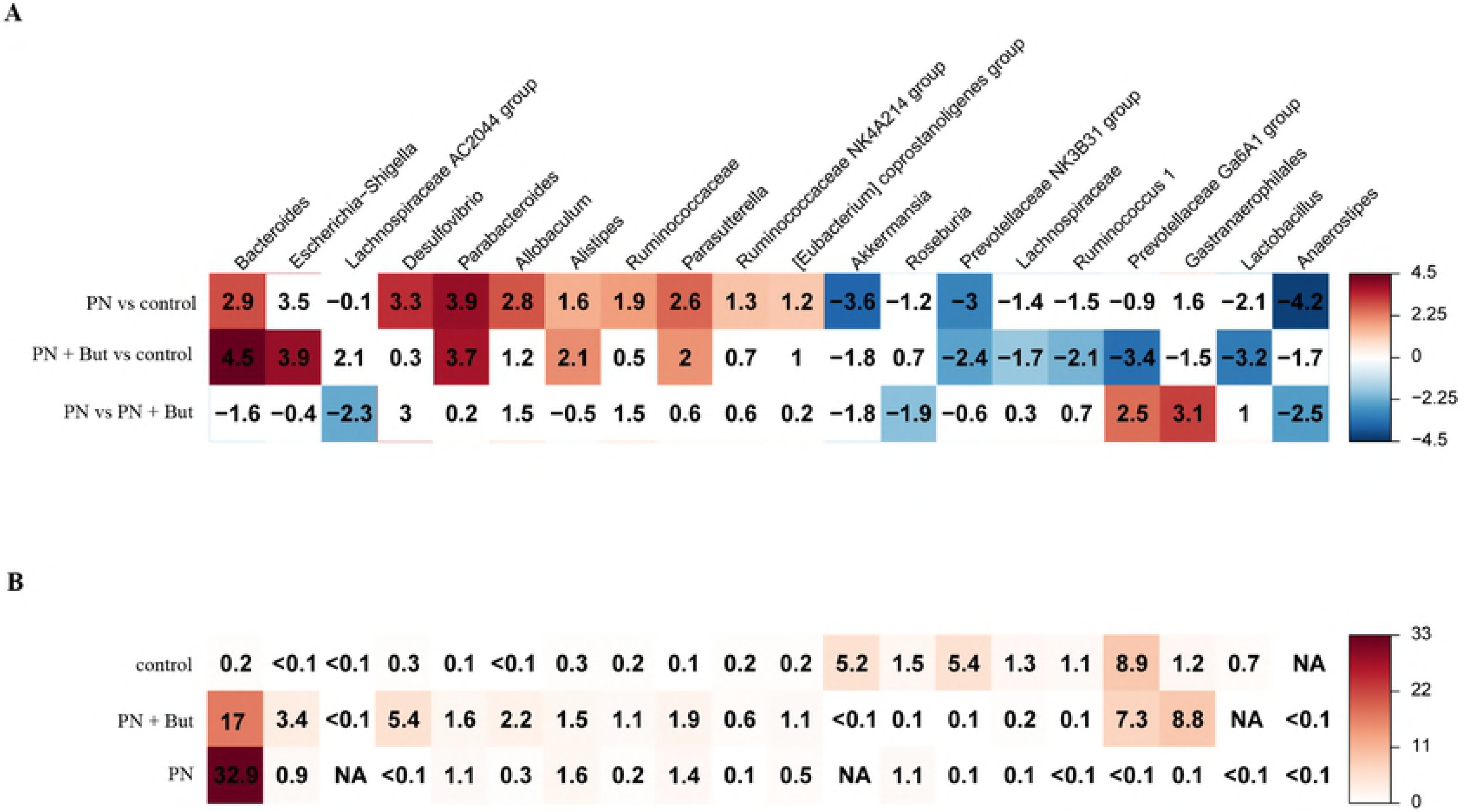
Heat map showing the fold change (A) and the abundance (%) (B) of genera that were differently (p<0.05) represented in at least two groups. Positive values correspond with an increase and negative values with a decrease in the first group compared with the second group. Shades of blue represents fold change decrease while shades of brown represent fold change increase. Uncoloured fields are not significant at p<0.05.

## Discussion

### Butyrate and non-immune defence systems

In the intestine, the basic line of defence (independent of immune cells) consists of a tight attachment of epithelial cells mediated by tight junction proteins, a mucin layer secreted by GCs, host defence peptides produced by Paneth cells, and enterocyte products like RegIIIγ and Muc3. All of these factors prevent bacteria from coming into contact with the sub-epithelial layer and thus inducing the inflammatory response. PN administration disturbs these systems (6, 7, 10, 30-33), resulting in the increased exposure of antigens to the immune system, increased intestinal permeability, and the establishment of pro-inflammatory status in the intestine.

Although there is abundant evidence (obtained both *in vitro* and *in vivo*) that butyrate affects all components of this defence system, the mechanism is not yet fully understood and controversies remain. Muc 2, secreted by GCs, is the major structural component of the intestinal mucus. Muc 3 is a transmembrane mucin produced by enterocytes and the major component of glycocalyx, which plays an active role in the intestinal mucosal defence (34). Studies published thus far have only focused on the effect of butyrate on mucin production when administered per rectum or in cell lines *in vitro*; furthermore, these results are rather inconsistent (35, 36). Gaudier (37) reported that, *in vitro,* butyrate grossly stimulated Muc2 expression but only in a glucose-deprived medium, while the effect of butyrate was dose-dependent and inhibitory at higher concentrations. These findings indicate that the effect of butyrate on mucus formation is context-dependent. The stimulatory effect of butyrate on host defence peptides and tight junction protein expression has been proved both *in vitro* and after dietary supplementation *in vivo* (38-42). Nevertheless, to our knowledge, no study has evaluated the effect of butyrate administered parenterally. Our data show that supplementation of a PN mixture with butyrate at a concentration within physiological limits (9 mM) upregulates the expression of all components of the non-immune defence – including mucins, host defence peptides and tight junction proteins in the ileum – while also improving intestinal permeability. We conclude that enforcement of the intestinal barrier may represent one of the beneficial effects of i.v. butyrate in the context of total dependence on PN and the absence of enteral nutrition and/or butyrate producers.

### Butyrate and immune functions

Total dependence on PN in critically ill patients is accompanied by decreased immune responsiveness, reduced gut-associated lymphoid tissue (GALT) mass, diminished IgA secretion, and increased risk of generalised sepsis (43). Nevertheless, it seems that the main factor responsible for immune dysfunction in PN-dependent patients is not PN administration itself, but the lack of enteral feeding (1). One consequence of the absence of enterally provided nutrients is low SCFA content in the gut. SCFA and, in particular, butyrate have been shown to influence immune cells towards anti-inflammatory and tolerogenic phenotypes (44) and to induce the differentiation of Foxp3+ Treg lymphocytes (45). In mice, an SCFA mixture administered *per os* increased the numbers of IgA-secreting lamina propria B cells, IgA expression or levels of secreted IgA in various compartments of the intestine, and IgA and IgG levels in the blood circulation (46). To our knowledge, only one study has focused on the effect of butyrate when added to PN on GALT. In mice, butyrate partially restored a PN-induced drop in lymphocyte numbers in Peyer’s patches and intestinal IgA levels (47). In our study, butyrate supplementation was associated with an increase in CD4+ lymphocyte numbers, and an increase in the CD4+/CD8+ ratio in MLN. Rather surprisingly, we observed an increase of Tregs in MLN of rats administered a PN mixture without butyrate, while the addition of butyrate resulted in a decrease in Tregs percentage to the control level. Treg cells expressing transcription factor Foxp3 are believed to play a key role in limiting inflammatory responses in the intestine (48), as they inhibit bystander T-cell activation either by a contact-dependent mechanism or through soluble factors (49). Paradoxically, Foxp3+ Tregs are more common in the inflamed intestinal mucosa of IBD patients, leading to a reciprocal drop in circulating Treg frequency in the peripheral blood; this likely reflects sequestration of these cells to the site of inflammation (50, 51). In a rat sepsis model, the pro-survival treatment was associated with a decrease in spleen Tregs (52) and in septic patients the persistence of elevated Treg indicated poor outcomes (53). We hypothesise that in our experimental setting decreased Treg frequency in MLN in the PN+But group reflects the lower inflammatory status of the intestinal epithelium, thus reducing the need to produce an anti-inflammatory response.

IgA production by plasmatic B cells in the submucosal layer is regulated by Th1 and Th2 cytokines produced by different T-helper subpopulations. While Th1 cytokines (IFNγ) downregulate IgA production, Th2 cytokines (IL-4, IL-5, IL-6, and IL-10) stimulate it (54). Hanna (55) reported that PN depressed both IL-4 and IL-10 levels in small intestine homogenates but that IFNγ levels remained unchanged, resulting in an imbalance between pro-/anti-IgA-regulating cytokines and a subsequent reduction in IgA production. Our data confirm this observation concerning the effect of non-supplemented PN. Butyrate supplementation resulted in increases in IL-4 and IL-10 expression to control levels and the near normalisation of IgA expression. These data suggest that intraepithelial Th2 helpers are one of the targets of butyrate and that butyrate supplementation may restore the PN-induced cytokine imbalance.

### Butyrate and the microbiota

The gut microbiome in animal models of PN is characterised by a significant shift in microbiota composition, particularly a loss of *Firmicutes* and an enrichment of *Bacteroidetes* and *Proteobacteria* (3, 7). In our study, we observed a shift towards an unfavourable microbiota composition, particularly an enrichment of *Proteobacteria* and the reduction of bacteria involved in butyrate production (*Lactobacillaceae* or *Lachnospiraceae*) in both PN-dependent groups. While the abundance of butyrate producers was not affected by butyrate supplementation we observed a trend, albeit not statistically significant, towards *Proteobacteria* reduction in butyrate-administered animals. Interestingly, butyrate supplementation (but not enteral deprivation/PN administration alone) was associated with a tendency to the loss of diversity.

Although the effects of dietary fibre on the gut microbiota have been described elsewhere (56), information concerning the direct effect of butyrate on the gut microbiota is scarce. Dietary butyrate was reported to reduce coliform bacteria (57) and to increase the abundance of *Lactobacillus* (42, 58) and butyrate producers *Blautia* and *Anaerostipes* (42). We observed no radical effect of *i.v.* butyrate, as it did not attenuate deregulation of the main contributors to PN-induced dysbiosis. Nevertheless, butyrate supplementation has been associated with an increased abundance of several potentially beneficial genera (*Anaerostipes, Roseburia, Lachnospiraceae AC2044 group*), a decreased abundance in the opportunistic human pathogen *Desulfovibrio* (59) and a trend towards attenuation in *Proteobacteria* dominance. We suggest that this subtle shift in microbiota composition may contribute, along with other mechanisms, to the overall beneficial effect of butyrate.

### Conclusion

We report that supplementation of a PN mixture with butyrate resulted in a significant enhancement of gut defence systems, i.e. increased expression of mucins, tight junction proteins and host defence peptides, and improvement of PN-induced aggravation of intestinal permeability. Lack of enteral nutrition and/or PN administration led to a shift in caecal microbiota composition. Although butyrate did not reverse the altered expression of most taxa, it did influence the abundance of several potentially beneficial or pathogenic genera what might contribute to its overall advantageous effect. We conclude that supplementation of a PN mixture with butyrate may represent a prospective therapeutic approach for mitigating the adverse effects of parenteral nutrition.

## Supporting information

Table S1. Composition of parenteral nutrition mixture

Table S2. Statistical analysis of sequencing data

## Author Contributions

**Conceptualization:** Monika Cahova

**Data curation:** Petra Videnska, Zuzana Jirsova, Monika Cahova

**Funding acquisition:** Monika Cahova

**Investigation:** Zuzana Jirsova, Marie Heczkova, Helena Dankova, Hana Vespalcova, Lenka Micenkova, Lenka Bartonova, Alena Lodererova, Alena Sekerkova

**Validation:** Lucia Prefertusová, Hana Malinska Eva Sticova, Petra Videnska

**Visualization:** Helena Dankova, Hana Vespalcova,

**Writing ± original draft:** Zuzana Jirsova, Petra Videnska, Marie Heczkova, Monika Cahova

**Writing ± review & editing:** Zuzana Jirsova, Petra Videnska, Monika Cahova

